# A Comprehensive DNA Methylome BodyMap across 12 Organs/Tissues from Spaceflight Mice

**DOI:** 10.64898/2026.07.08.737015

**Authors:** Zhong Chen, Chirag Nepal, Xiao Wen Mao, Feng Zeng, Michael Pecaut, Marjan Boerma, Charles Wang

## Abstract

Spaceflight imposes unique physiological stresses on mammals, including microgravity and cosmic radiation, which drive complex molecular adaptations. However, the systemic and temporal dynamics of space-induced epigenetic regulation remain poorly understood. We constructed a comprehensive DNA methylome BodyMap across 12 organs or tissues from mice exposed to long-duration spaceflight across three time points using Reduced Representation Bisulfite Sequencing (RRBS). We also performed RNA-seq for five organs and integrated with DNA methylome. We mapped the methylome and transcriptome landscapes and found that spaceflight induces limited but highly tissue-specific differentially methylated CpGs (DMCs). Most spaceflight-induced methylation changes were reverted toward baseline within one to six months of post-flight. Functional enrichment analysis of DMCs highlighted metabolic and mitochondrial dysregulation commonly across organs, while developmental responses in immune, reproductive, and structural tissues were tissue-specific. Transcriptome data revealed that spaceflight suppressed immune and increased inflammatory responses at the multi-organ level, triggering a phenomenon resembling aging. Our study provides a comprehensive DNA methylome BodyMap across 12 organs/tissues in spaceflight mice, elucidating the tissue specificity of epigenetic changes. These insights are essential for developing biomarkers and countermeasures to safeguard astronaut health during extended missions.

## Introduction

Spaceflight exposes biological systems to multiple environmental stressors, such as microgravity, cosmic radiation and circadian rhythms, that profoundly challenge mammalian physiology^1,2^. These conditions trigger complex molecular adaptations in human homeostasis, function, and health, manifested by the loss of muscle mass and bone density, as well as immune dysregulation^3^. As humans embark on extended missions to the Moon and Mars, understanding the molecular mechanisms underlying these physiological changes is essential for developing effective countermeasures. Gene expression and epigenetic regulation are key mediators of cellular adaptation to environmental stress, yet how these processes respond to spaceflight remains incompletely understood.

DNA methylation plays an essential role in mammalian organ development, cellular identity, and environmental adaptation^4,5^. Spaceflight-induced DNA methylation changes are generally subtle, affecting limited CpGs sites^1,3^ and often reverting to baseline after returning to Earth^3^. The functional relationship between DNA and gene expression during spaceflight remains unclear. Analysis of NASA Rodent Research (RR-1 and RR-3) GeneLab data^1^ and JAXA studies revealed minimal promoter methylation changes weakly correlated with gene expression^6^. Spaceflight studies in mice retinas identified a few hundred CpGs whose methylation changes were associated with altered gene expression^7^, whereas rice germ cells exhibited hypermethylation without corresponding transcriptional changes^8^. Additional studies indicate that space-induced epigenetic alterations may be inherited^9^. Microgravity simulation revealed hypomethylation and mutations in human lymphocytes^10^. Most differentially methylated regions in cultured human lymphoblastoid cells under simulated gravity were hypomethylated and linked to oxidative stress response and carbohydrate metabolism^11^. Induced genome-wide methylation changes on hindlimb experiments were mostly hypermethylated on CpG shores^12^, which may be due to increased expression of genes, such as Dnmt1, Dnmt3a, and MecP2^13^, that regulate DNA methylation.

Despite these findings, the systemic and temporal dynamics of space-induced epigenetic regulation remains largely unexplored. Comprehensive, multi-tissue profiling is required to elucidate how prolonged spaceflight influences DNA methylation and transcriptional programs across organ systems. To address this, we analyzed DNA methylation across 12 mouse tissues and transcriptomes from 5 organs collected during NASA’s Rodent Research-18 mission, generating a multi-organ DNA methylome atlas of long-duration spaceflight. Our goal was to generate a comprehensive DNA methylome body map during long-term spaceflight, spaceflight-induced DNA methylation changes, and define the reversibility or persistence of epigenetic changes. We aimed to uncover global trends linking DNA methylation changes with gene expression dynamics. Our integrative, multiorgan, multi timepoint approach offered a comprehensive view of how long-duration spaceflight reshapes the epigenome and transcriptome across organs/tissues, providing biomarkers and mechanistic insights essential for astronaut health monitoring and countermeasure development.

## Results

### Study design of spaceflight induced DNA methylome and transcriptomic landscape

We profiled the DNA methylome and transcriptome of 16-week-old male C57BL/6 mice exposed to spaceflight. Animals were assigned to three groups under two conditions (ground control and spaceflight) (Fig. 1). The first group was euthanized after 75 days aboard the International Space Station (ISS) on the rodent research-18 (RR-18) mission and immediately froze at -80°C at ISS (FZ samples). The second and third groups were flown aboard the ISS as part of the RR-18 over 35 days and returned back to Earth alive. The second group was euthanized within 24 hours after the live animal returned (LAR_acute). The third group was euthanized 3 months after returning to Earth (LAR_3M). Matched ground controls (GC) were euthanized three days later, and tissues collected at each corresponding time point. After dissection, tissues were frozen at -80°C until RNA/DNA extraction.

**Figure 1.**
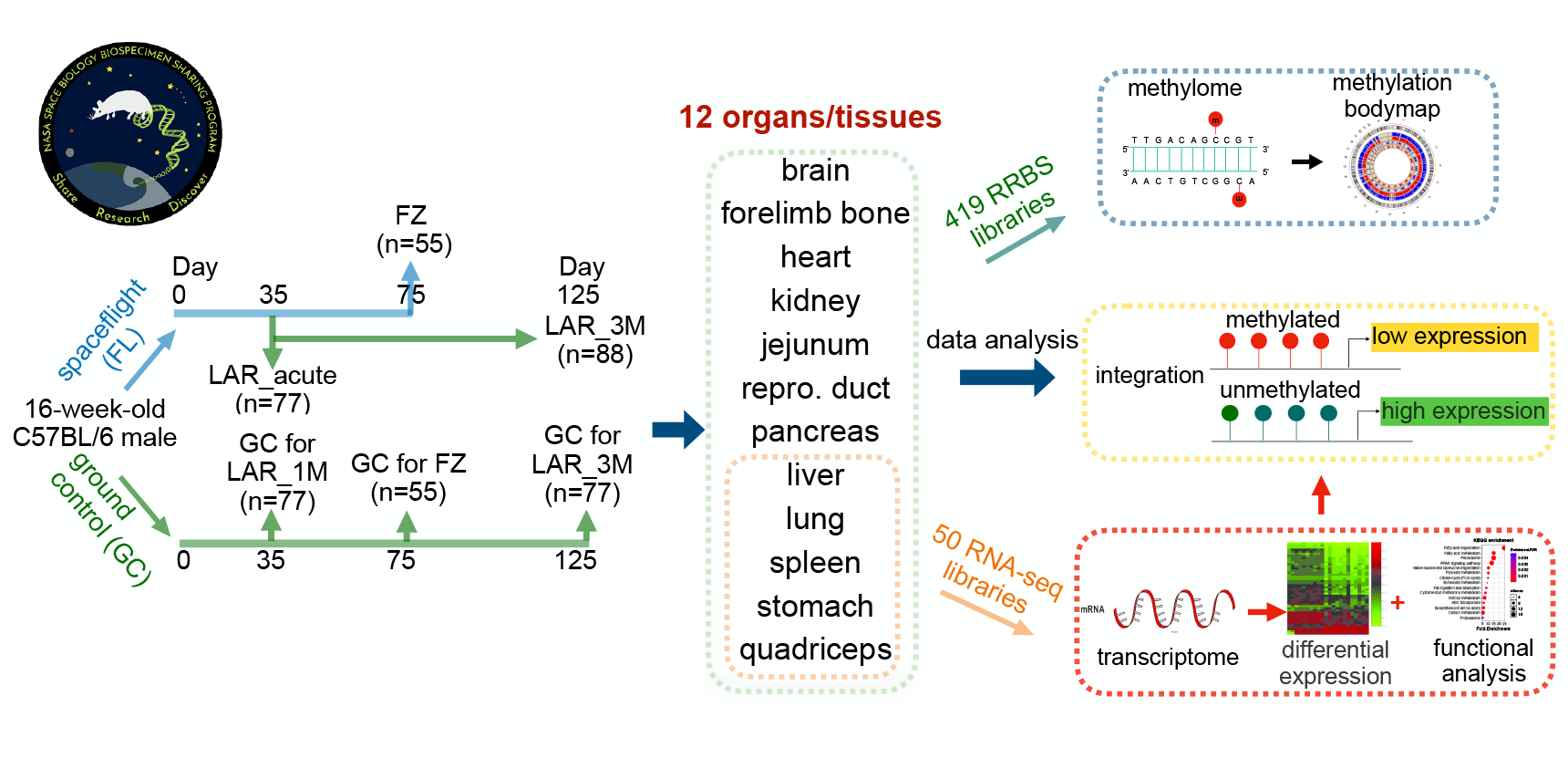
Schematic overview of study design. 16-week-old male C57BL/6 mice were used in this study. One group was sent to the International Space Station (ISS) and another group was kept as ground control in the same vivarium. The timeline illustrates the duration mice were exposed to spaceflight and the recovery time after returning to Earth. Animals were euthanized at three time points: euthanized and frozen at ISS after 75 days (FZ); returned to Earth alive and dissected within 24 hours (Liver animal return 1day; LAR_acute) and recovered for 3 months (LAR_3M). All eleven organs/tissues were used for DNA methylation, and five organs/tissues were used for transcriptome analysis.

To generate DNA methylome maps, we constructed 419 Reduced Representation Bisulfite Sequencing (RRBS) libraries from twelve organs/tissues across all three time points with 5-8 mice in each of the spaceflight and ground control group (**Fig. 1)**. We obtained an average of 3.5 million CpG sites mapped to the NCBI mouse GRCm38 genome (**Table S1, Fig. S1a-c**). To generate consensus CpGs for each tissue, we ensured that CpGs were detected in all samples and required to have a minimum of 5 reads in at least 2 samples, which resulted in an average of 1.41 million CpGs per tissues/organs. For downstream analysis, we ensured that CpG sites were detected in all samples across a given tissue. For transcriptomics data, we constructed 50 total RNA libraries from five organs/tissues (liver, lung, spleen, stomach, and quadriceps). Each sample was sequenced to an average of 30 million 150x2 paired-end reads. Sequence reads were mapped to the NCBI mouse GRCm38 genome with over 90% of unique mapping rate (**Fig. S1d**). Mapped methylome and transcriptomic data were systematically analyzed to identify spaceflight induced changes.

### DNA methylome landscape of twelve mouse organs/tissues between spaceflight and Earth conditions

Different genomic regions exhibit distinct methylation levels and patterns^4,14^, prompting us to analyze global DNA methylation levels across twelve organs/tissues in ground and spaceflight conditions. For each tissue, we computed mean CpG methylation levels (0-1) across all replicates from ground and spaceflight conditions. Using bone as an example, CpGs methylation exhibited a bimodal distribution with most CpG either unmethylated (beta value < 0.1) or fully methylated (beta value > 0.9) in both ground and spaceflight (FZ) samples (**Fig. 2a-b**). This bimodal pattern was consistent across other tissues and time points between ground and spaceflight (**Fig. 2c-d**) and also in ground control and spaceflight LAR_acute groups (**Fig. S2a-b**). Overall, spaceflight did not globally alter DNA methylation (**Fig. 2c-d**), suggesting that stable epigenetic states established during development are maintained to ensure organ identity and function^15^. To examine methylation levels across different genomic regions, we plotted profiles over gene bodies, promoter CpG islands (CGIs), and intragenic CGIs. Transcription start sites (TSSs) exhibited the lowest methylation, which increased gradually from TSSs toward both ends in ground and spaceflight conditions (**Fig. 2e**). Gene bodies had the highest methylation relative to 5’- and 3’-flanking regions. Promoter CGIs, which largely overlap with CGIs^16^ were hypomethylated (**Fig. 2f**), consistent with canonical low methylation in normal tissues^4^. In contrast, intragenic CGIs, which generally lack TSSs^17^ had higher methylation comparable to its flanking regions (**Fig. 2g**).

**Figure 2.**
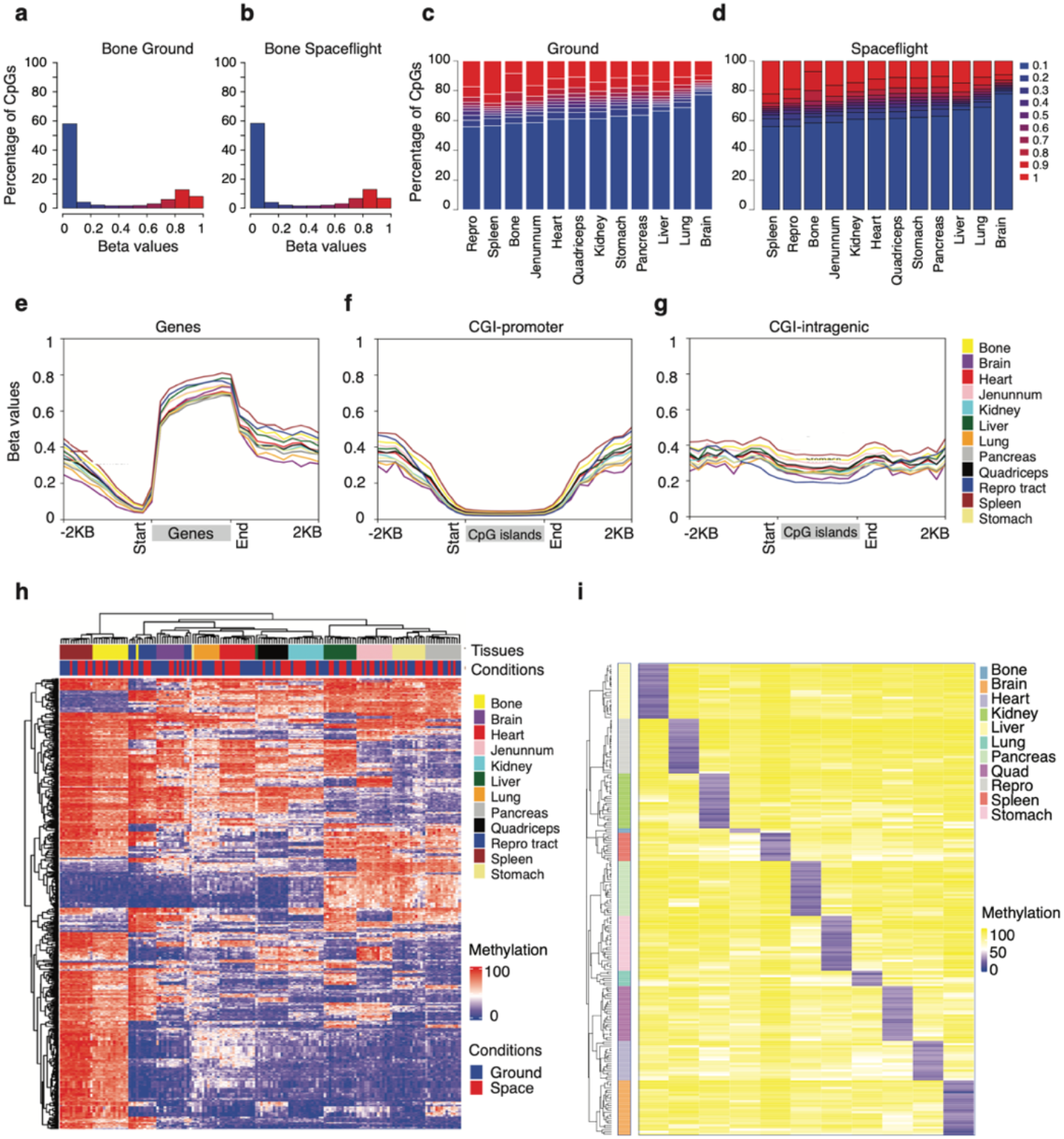
DNA methylation landscape across conditions and organs/tissues. (**a-b**) Distribution of mean beta values of all CpGs from bone tissues in ground (**a**) and spaceflight (**b**) conditions. X-axis represents beta values, which are divided into ten bins. Y-axis represents the number of CpGs in percentage. (**c-d**) Distribution of mean beta values of all CpGs from all tissues in ground (c) and spaceflight (**d**) conditions. Y-axis represents the percentage of CpGs in each beta value bin. (**e-g**) Distribution of mean beta values of all CpGs across all genes (e), genes with promoter CpG islands (CGIs) (**f**) and genes with intragenic CGIs (g). (**h**) Heatmap showing the hierarchical clustering of highly variable CpGs that cluster samples according to tissue type. (**i**) Heatmap shows blocks of tissue-specific hypomethylated CpGs that are hypermethylated in other tissues.

DNA methylations reveal tissue-specific methylation patterns^18,19^, thus we asked whether organ/tissue methylation distinguishes ground tissues from spaceflight conditions. To this end, we calculated the variance of all CpGs across all tissues from ground and spaceflight (methods). We performed hierarchical clustering using the top one percentile of CpGs and observed that samples were clustered by tissue types, with no clear separation between ground and spaceflight within tissues (**Fig. 2h**). We then identified the most variable (>98 percentiles) CpGs across all organs/tissues separately from ground and spaceflight samples and observed high (>90%) overlap (**Fig. S2c**). This suggests that tissue-specific methylation is the dominant source of variation in both conditions, and spaceflight disturbs global DNA methylation at a small scale after 35-day duration (equivalent to 4 human years). Human cell types have distinct unmethylated CpG blocks that are methylated in other tissues^19^, thus we sought to identify such patterns in mice. We identified tissue-specific hypomethylated CpG in each organ/tissue (**Fig. 2i**). Uniquely hypomethylated regions (average methylation < 15% in a given tissue but >70% in other tissues) can serve as tissue-specific biomarkers. This suggests that spaceflight-induced methylation changes likely affect non-tissue-specific CpGs, or they induce subtle methylation changes masked by tissue-specific variability.

### Spaceflight-induced differential methylome across different organs/tissues

As the above unbiased clustering of top variable CpGs revealed no global separation between ground and spaceflight samples, we hypothesized that spaceflight-induced methylation changes might be subtle. To investigate this, we identified differentially methylated CpG sites (DMCs) between spaceflight and ground controls for each organ/tissue using linear regression implemented in the limma package^20^. DMCs were defined at the threshold of p-value < 0.001 and absolute methylation difference (beta value (>10%). Across tissues in the FZ group, we identified an average of ∼4,000 DMCs (**Fig. 3a; Table S2**), representing <0.5% of analyzed CpGs, indicating limited and tissue-specific methylation alterations. In the LAR_acute group, the number of DMCs in most tissues were generally lower than that found in FZ group, except for brain and spleen (**Fig. 3b; Table S3**), suggesting that many early spaceflight-induced epigenetic changes had reverted after one month of recovery. In the LAR_3M group, most tissues showed increased number of DMCs compared to FZ group, with bone exhibiting a particularly large number of DMCs (**Fig. 3c; Table S4**), possibly indicating long-term compensatory changes or delayed secondary epigenetic remodeling.

**Figure 3.**
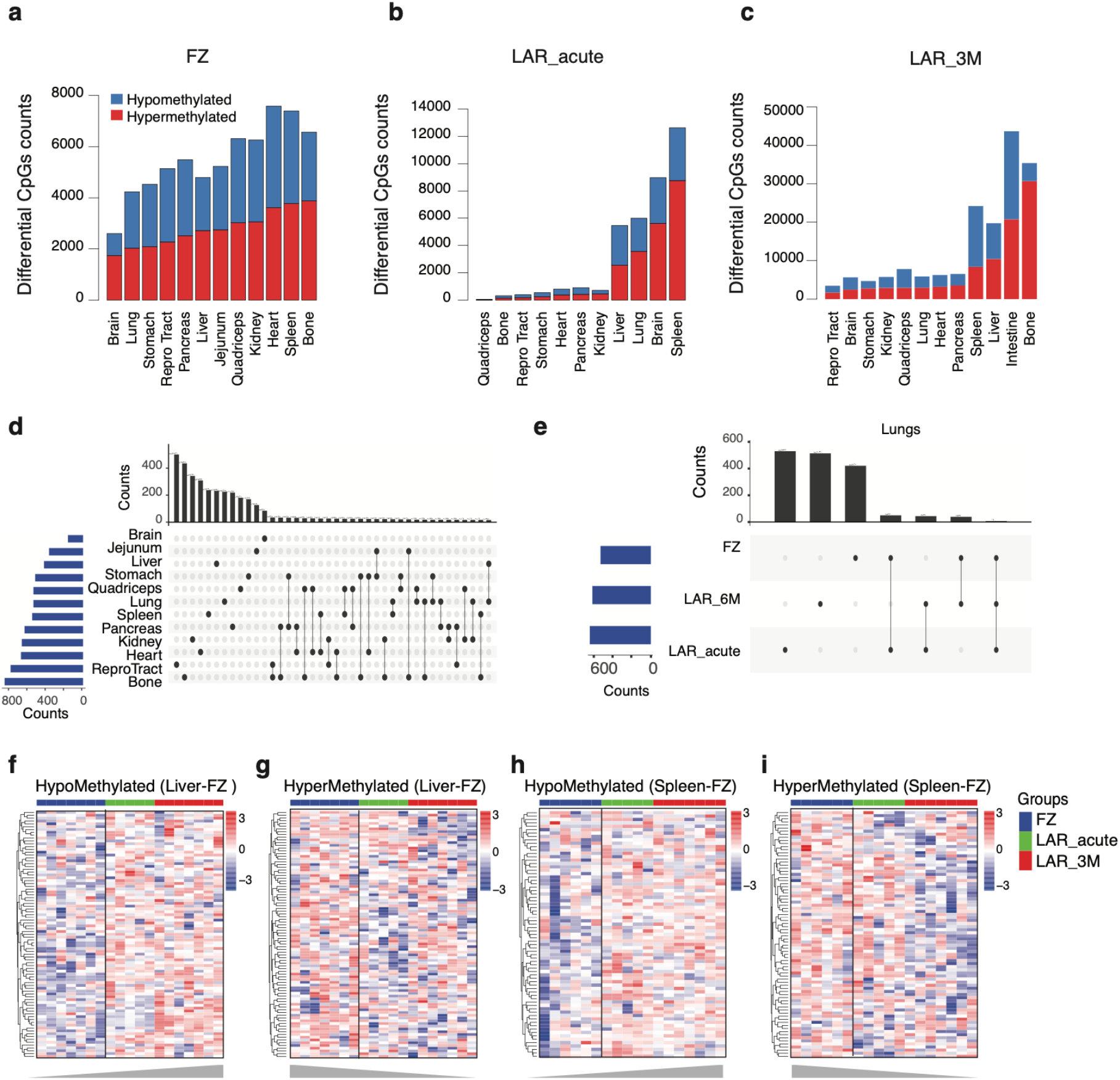
Landscape of spaceflight induced differentially methylated CpGs. **(a-c)** The number of spaceflight induced differentially methylated CpGs across different tissues in flight group (FZ) (a) and two recovery samples at LAR_acute (b), and LAR_3M (c). (**d**) Overlap of differentially methylated CpGs between different organs/tissues in FZ group. (**e**) Overlap of differentially methylated CpGs from lung tissues during flight (FZ) and two recovery time points (LAR_acute and LAR_3M). The majority of differentially methylated CpGs are unique to each time point. (**f-g**) Heatmaps of differential hypomethylated CpGs (f) and hypermethylated CpGs (g) identified in liver during flight (FZ) and their methylation levels in corresponding recovery samples at LAR_acute and LAR_3M. Methylation values are row-scaled. (**h-i**) Heatmaps of differential hypomethylated CpGs (h) and hypermethylated CpGs (i) identified in spleen during flight (FZ) and their methylation levels in corresponding recovery samples at LAR_acute and LAR_3M. Methylation values are row-scaled.

Differentially methylated CpGs identified in the FZ group exhibited minimal overlap across tissues (**Fig. 3d**), a pattern that was also observed in the LAR_acute and LAR_3M groups (**Fig. S 3a-b**), suggesting that spaceflight-induced methylation changes are highly tissue-specific. Likewise, comparison of lung DMCs across the FZ, LAR_acute, and LAR_3M groups revealed little overlap (**Fig. 3e**), indicating that most spaceflight-induced methylation changes likely reversed after returning to Earth. Consistent patterns were observed across other tissues (**Fig. S3c-f**). To examine these dynamics, we tracked methylation levels of FZ DMCs across recovery (LAR_acute and LAR_3M) timepoints. In the liver, FZ hypomethylated DMCs showed progressive remethylation in LAR_acute and LAR_3M (**Fig. 3f**), while hypermethylated DMCs showed progressive demethylation (**Fig. 3g**), consistent with reversion toward baseline levels. Similar patterns of recovery were observed in spleen DMCs (**Fig. 3h-i**) and brain DMCs (**Fig. S3g-h**), where both hyper- and hypomethylated sites trended back to preflight levels. To test whether these trends reflected recovery from spaceflight rather than temporal variation, we evaluated the same DMCs in ground controls across FZ, LAR_acute, and LAR_3M groups. In contrast to the flight groups, ground controls showed no evidence of systematic reversal in either the liver, brain or spleen (**Fig. S3i-n**). These results suggest that most spaceflight-induced methylation changes are reversible and resolved after returning to Earth.

### Spaceflight induced DNA methylation changes are linked to distinct biological processes and transcription factor motifs

To elucidate the biological processes associated with spaceflight induced DMCs in FZ group, we analyzed Gene Ontology (GO) using clusterProfiler^21^. Proximal DMCs near genes can regulate gene expression^4^, thus we associate DMCs within 10KB upstream and downstream of genes start and end. We observed that some biological processes, particularly those metabolic processes related to vitamin metabolism, terpenoid and retinoic acid biosynthesis, and mitochondrial transcription, were enriched across tissues (**Fig. 4a, Fig. S4a-b**), which is consistent with previous observation that highlighted mitochondrial dysregulation as a major consequence of spaceflight^1^. Enriched GO terms in cellular stress responses (toxic substance response, interferon response) exhibited a distinct pattern across tissues, suggesting tissue-specific epigenetic adaptation to spaceflight conditions. In contrast, developmental and structural processes, including skeletal muscle development, skeletal system morphogenesis, and embryonic organ morphogenesis, were broadly downregulated (blue), especially in muscle, bone, and heart tissues. Immune and reproductive tissues were enriched for cellular response to chemical stress, cell cycle regulation, and cellular senescence that exhibited more tissue-specific patterns. Several GO terms related to cellular homeostasis, including cell cycle G1/S phase transition, regulation of cellular senescence and T cell mediated cytotoxicity were enriched in immune-related tissues (spleen, bone). The hierarchical clustering of enriched GO terms revealed distinct patterns, with metabolically active tissues (liver, pancreas, kidney) clustering separately from structural tissues (bone, muscle), suggesting coordinated epigenetic responses within functionally related tissue types during spaceflight exposure. Together, these results indicate that spaceflight induced DMCs highlight tissue-specific metabolic and structural adaptations, offering insight into the molecular mechanisms underlying physiological adaptation to microgravity and space radiation exposure.

**Figure 4.**
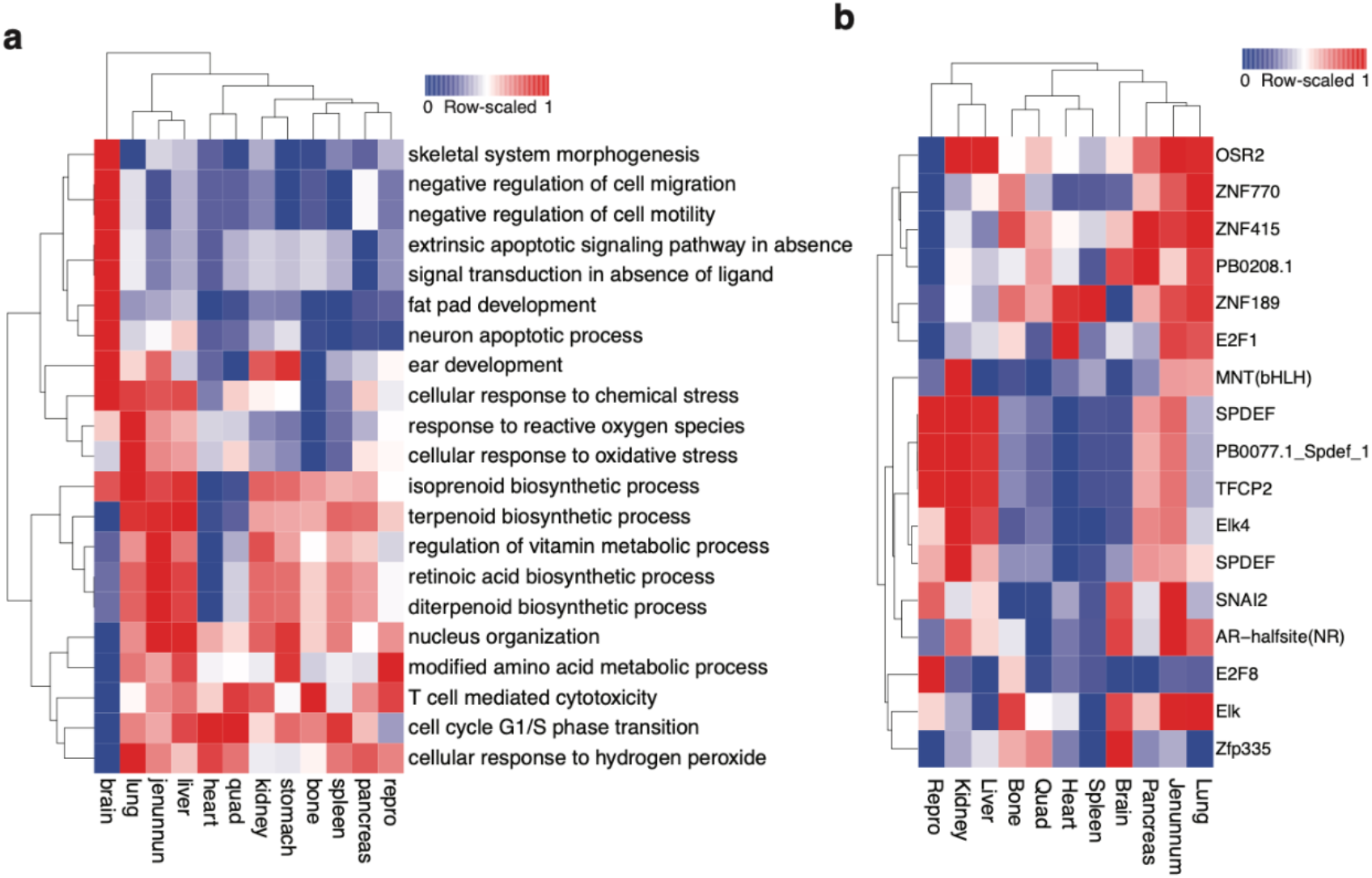
Enrichment of gene ontology and transcription factor motifs associated with differentially methylated CpGs (DMCs) from FZ group. (a) Heatmap shows enriched gene ontology terms on genes proximal to DMCs. P-values are scaled across rows. (b) Heatmap shows transcription factors motifs enriched on DMCs across various tissues. Selected motifs are enriched (p-value < 10e-11) at least in one tissue/organ. Heatmap shows the fold-enrichment of target versus background sequence, and it is row-scaled in the range of 0-1.

To identify enriched transcription factor (TF) associated with DMCs, we extended 500 bases around DMCs and performed enrichment analysis using HOMER^22^. We identified several TF motifs were enriched in a tissue-specific manner (**Fig. 4b**). Motifs for OSR2 and zinc finger proteins (ZNF770, ZNF415, ZNF189) were enriched in bone, heart, and reproductive tissues, reflecting the roles of those TFs in cell differentiation and development. In contrast, E2F1, E2F8, and MNT motifs, which are key regulators of the cell cycle, were enriched in pancreas and spleen. Neural and immune tissues showed preferential enrichment for ETS family motifs (Elk1 and Elk4), while SPDEF and SNAI2, involved in epithelial identity and migration, were enriched in lung and kidney. Notably, the androgen receptor (AR) motif was uniquely enriched in reproductive tissues, manifesting its role in male sexual development. Together, these findings demonstrate that tissue-specific DNA methylation patterns are closely tied to the regulation of key biological pathways and the selective recruitment of transcription factors. Collectively, these findings suggest that spaceflight-induced DNA methylation changes may be coordinated by a distinct set of transcription factors, potentially mediating tissue-specific adaptation to the space environment.

### Spaceflight-induced transcriptomic alterations across organs and tissues

To understand the spaceflight-induced transcriptomic alterations, we profiled the transcriptomes of 5 organs/tissues from the FZ group and their matched ground controls. Principal component analysis (PCA) revealed that spaceflight markedly altered the transcriptomes in all five organs (**Fig. S5a-e**). Among all organs/tissues, the liver showed the strongest impact, as the PC1 clearly separated spaceflight samples from the ground controls. We then identified the differentially expressed genes (DEGs) in each organ (**Fig. 5a-e**), using the cutoff of FDR<0.01 and |log2FC|>0.5. Liver had the highest number of DEGs (404 upregulated and 322 downregulated genes) compared to about 300 DEGs in lung, spleen, stomach, and quadriceps, respectively (**Fig. 5a-f, Table S5)**. The DEGs across five organs/tissues had low overlap indicating that the majority of the DEGs are tissue-specific, where liver and lung shared the largest number of DEGs (**Fig. 5f**). GO analysis showed that fatty acid and lipid metabolism were significantly enriched in the liver and stomach from their individual DEGs (**Fig. 5g**) or common DEGs (**Fig. S6d**). Positive z-scores (**Fig. S5f, 5i**) and increased transcription levels (**Fig. S6a**) indicated a synergistic upregulation of fatty acid and lipid oxidation in spaceflight liver and stomach. Liver, lung, and spleen showed similar enrichment in interferon signaling and antigen presentation with strong negative z-score (**Fig. 5g, Fig. S5f-h, Fig. S6e, 6f**) and suppressed gene expression, indicating that spaceflight trigged the immunodeficiency in multiple organs and leading to reduced antiviral ability (**Fig. S6b, 6c**). The top enriched canonical pathway in quadriceps was the response to unfolded protein (**Fig. S5j, Table S6**), consistent with the notion that spaceflight induces muscle atrophy. It is worth noting that the PPAR signaling pathway, which regulates metabolic homeostasis and function in multiple organs, was strongly enriched (Foldchange > 40) in stomach and liver common DEGs (**Fig. S6d**). Many of the genes related to the inmate immune system were significantly suppressed (**Fig. S6g**). The biological functions of the proteins encoded by those 9 genes covers many aspects of inmate immune response, including viral RNA recognition (*Dhx58*), viral RNA degradation (*Oas2*, and *Oas1a*), antigen processing and presentation (*Psmb10*), as well as B cell development (*Bst2*). Our data strongly indicated a comprised inmate immune system in multiple organs of mice flown at ISS, highlighting the adverse impact of spaceflight on immune system that were observed in other model animals and astronauts subjected to spaceflight stressors^1,23^. To understand whether promoter DNA methylation has a significant role in regulation of gene expression, we correlated methylation levels and gene expression levels. Overall, we observed a negative correlation between promoter methylation and gene expression (**Fig. 5h-i**), similar to NASA RR-1 and RR-3 GeneLab data^1^. Individual examples of genes *Clcn2* and *Ldha* depict a negative correlation between methylation levels and gene expression (**Fig. S5k-n**). As the majority of promoters had low methylation levels, which did not change in spaceflight, a low negative correlation was expected. In sum, we observed that spaceflight altered the expressions of genes involved in lipid oxidation and immune response in a concerted way in multiple organs, however those transcriptional changes were not directly regulated by spaceflight induced DNA methylation change.

**Figure 5.**
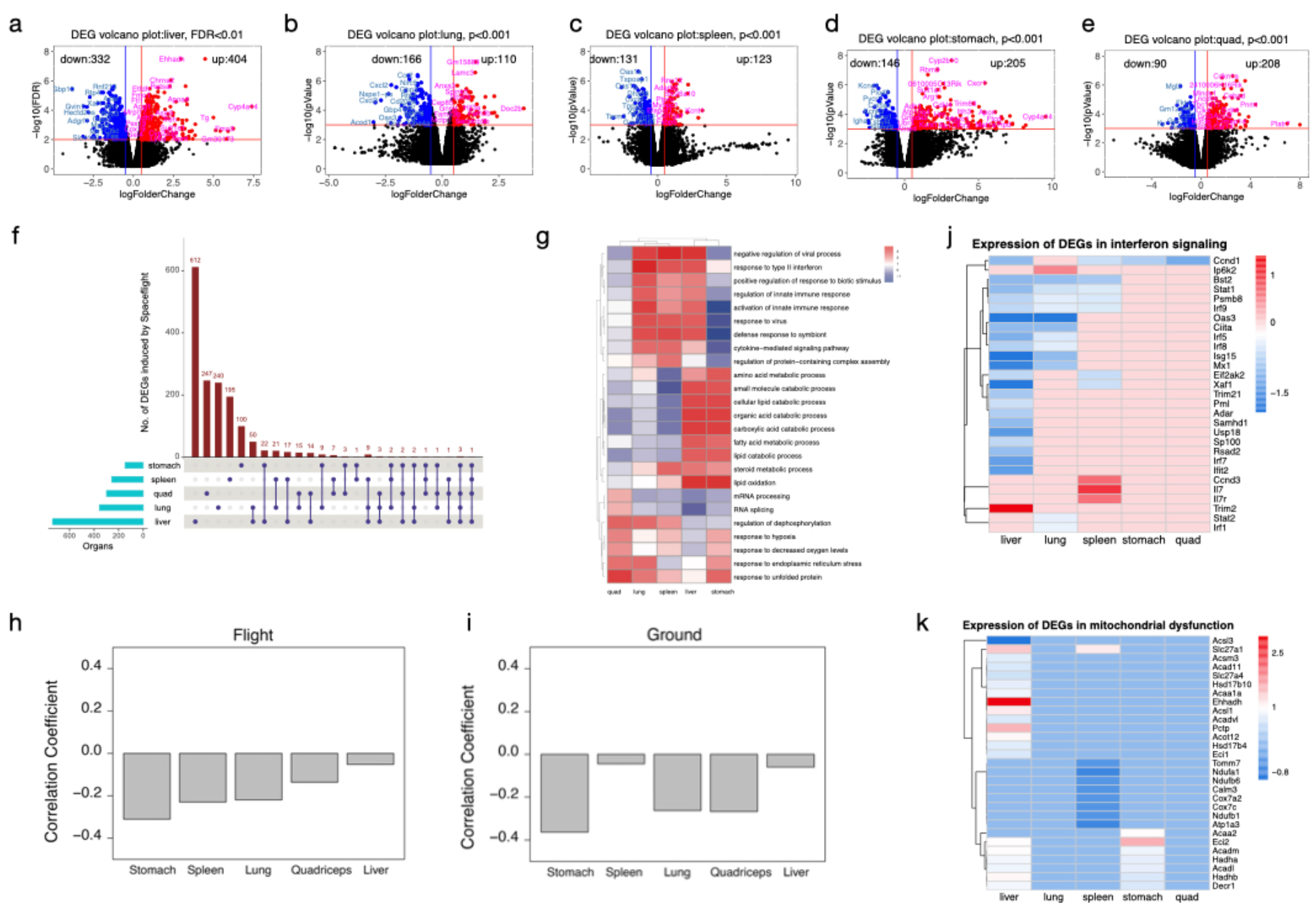
Spaceflight altered gene expression across multiple organs. **(a-e)** Volcano plots showing the differentially expressed genes (DEGs) induced by spaceflight in liver (a), lung (b), spleen (c), stomach (d), and quadriceps (e). Vertical lines showing the log2FC cutoff at -0.5 and 0.5, and the horizontal line shows the FDR or p value cutoff. (f) Upset Plot showing the number of DEGs in each organ and the number of overlapped DEGs. (g) Heatmaps showing enriched Gene Ontology biological processes on DEGs induced by spaceflight in each organ. Spaceflight elicited concordant responses to multiple organs. Enrichment scores are row-scaled. (h-i) Barplots showing the global correlation coefficient of all genes and their promoter methylation levels in flight (h) and in ground group (i). (j) Heatmap showing the fold-change of DEGs from interferon signaling pathway. Expression values are row-scaled. (k) Heatmap showing the fold-change of DEGs from mitochondrial dysfunction. Expression values are row-scaled.

### Spaceflight impacts hallmarks of aging

The majority of the common DEGs between liver and lung, liver and spleen were predominately involved in antiviral and immune deficiency (**Fig. 5j, Fig. S6d-f)**, and their expressions were concordantly suppressed under spaceflight conditions (**Fig. S6b,6c)**. For example, spaceflight significantly suppressed the expression of genes in interferon alpha/beta signaling (IFN-α and IFN-β are type I IFNs) and the activation of Th1/Th2 (T helper) (**Fig. 5j**). IFN-α and IFN-β are key components of the innate immune system, primarily involved in the antiviral response and IL-4 is a cytokine primarily produced by Th2 cells which is essential for inflammatory response, such as allergic reactions and asthma. It is known that aging is associated with reduced production of IFNs and dysregulation of inflammatory immune responses, a phenomenon known as “inflammaging”^23,24^. This view is supported by the transcriptomic changes observed in multiple organs under spaceflight in this study. The common upregulated DEGs in liver and stomach were enriched in fatty acid degradation in a synergistic manner (**Fig. 5k, Fig. S6a, Fig. S6d**). Elevated fatty acid beta-oxidation will increase the production of reactive oxygen species (ROS) in mitochondria. When ROS production overwhelms the cell’s antioxidant defenses, it will damage the cellular components like DNA, proteins, and lipids, manifesting many hallmarks of aging^25^. We also found that spaceflight impacts the muscle macromolecule biosynthesis and protein folding in the quadriceps of spaceflight mouse (**Fig. 5g, Fig. S5j**) that suggests muscle atrophy. Muscle atrophy is a phenomenon prevalent in the elderly population and also commonly recorded in astronauts^26^ and in space mission rodents^27,28^. Our result hinted that spaceflight triggers physiological deterioration resembling aging.

## Discussion

In this study, we present a comprehensive multi-organ mouse body map of DNA methylation and gene expression as well as spaceflight induced changes on DNA methylation and gene expression. This dataset, which integrates RRBS and RNA-seq data from multiple organs/tissues, represents one of the most extensive epigenomic and transcriptomic resources currently available from a long-duration NASA Rodent Research mission. By combining tissue-specific and shared molecular responses, this dataset can be exploited by space biology and mouse research communities to uncover regulatory principles governing adaptation to microgravity and space radiation. The integrative analysis pipeline and curated results also provide a foundation for hypothesis generation and testing in future studies, including those involving precision medicine countermeasures, immune modulation, and tissue-specific vulnerability assessments.

While the epigenome is sensitive to environmental factors, we observed a relatively modest number of differentially methylated CpGs (DMCs) across tissues, which is consistent with previous studies^1,3,7^. Several factors may account for this observation. First, DNA methylation is a relatively stable epigenetic mark, and short-term or mid-duration exposures (e.g., 30–90 days) may not yield extensive methylation reprogramming. Second, in adult tissues, most CpGs are either fully methylated or unmethylated and remain static in the absence of significant developmental cues or environmental disruption. Third, it is possible that spaceflight-induced transcriptional changes are primarily regulated via non-DNA methylation epigenetic mechanisms, such as histone modifications, noncoding RNAs, or 3D chromatin remodeling. Finally, the tissue-specific nature of the DMCs, as well as their proximity to regulatory regions rather than core promoters, may reflect a fine-tuning role of DNA methylation in orchestrating context-dependent gene regulation during spaceflight rather than wholesale reprogramming.

In contrast to the limited number of DMCs, we observed a robust transcriptional response to spaceflight, with hundreds of differentially expressed genes (DEGs) per organ. This discrepancy between DNA methylation and gene expression changes highlights the dynamic and multifactorial nature of transcriptomic regulation. Gene expression can rapidly respond to external stimuli through transcription factor activity, RNA stability modulation, and chromatin accessibility changes, all of which may be more acutely sensitive to the microgravity environment and radiation exposure than DNA methylation. This pattern of strong transcriptional responses and more stable epigenetic marks has been observed in previous NASA GeneLab rodent missions^1,3^ and underscores the importance of integrating multiple omics layers to capture the full scope of physiological adaptation to space.

One of the most consistent findings across both methylation and gene expression datasets was the alteration of mitochondrial and metabolic pathways, particularly in the liver, brain, and jejunum. GO enrichment of genes proximal to DMCs and DEGs highlighted pathways including fatty acid oxidation, PPAR signaling, retinoic acid metabolism, and mitochondrial transcription. The pronounced upregulation of key metabolic regulators such as *Ehhadh* and *Cyp4a14* further supports a shift in energy balance and mitochondrial stress. Mitochondrial dysfunction has been a recurring theme in spaceflight studies and is increasingly recognized as a central node linking oxidative stress, immune dysfunction, and altered tissue homeostasis. Our data further reinforce this concept and point to the liver and stomach as central hubs of metabolic adaptation under microgravity. Immune suppression and dysregulation are known consequences of spaceflight, and our multi-organ data corroborated this phenomenon. Downregulation of innate immune genes in the spleen and lung, including antiviral defense genes (e.g., *Oas1a, Oas2, Dhx58, Rnf21*, and *Gbp1*), indicates impaired immune readiness. The IPA enrichment of pathways such as NOD-like receptor signaling and primary immunodeficiency, particularly in genes shared between liver, lung, and spleen, suggests systemic immune suppression. Importantly, the observed changes in gene expression occurred in the absence of viral infection, implying that the altered immune landscape is likely driven by intrinsic factors such as microgravity-induced cellular stress or radiation-induced DNA damage. Spaceflight triggered clear downregulation of developmental and structural gene sets in muscle, bone, and heart, consistent with known atrophic effects of microgravity on musculoskeletal systems. In the quadriceps, DEGs were enriched for RNA processing, protein turnover, and cell cycle regulators, with notable induction of *Cdkn2a*, a gene associated with cellular senescence and muscle atrophy. These findings are aligned with previous rodent and human studies in space and support the hypothesis that epigenetic and transcriptional alterations contribute to the degradation of skeletal integrity in space environments.

Although most gene promoters maintained low and static methylation, a subset of genes with altered promoter methylation levels showed an inverse correlation with gene expression. This negative correlation, although modest in magnitude, is consistent with established paradigms and previous NASA RR studies. However, the relatively limited overlap between DMCs and DEGs suggests that promoter methylation is one of several layers of regulation influencing gene expression during spaceflight.

Many canonical pathways altered by spaceflight resemble the hallmarks of aging, like reduced immune response, mitochondrial dysfunction, and skeleton muscle atrophy. In this study, we found that spaceflight suppressed the macrophage activation, cytokine storm signaling, and the interferon alpha, beta, and gamma signaling in multiple organs, inducing an inflammatory status known as inflammaging (**Fig. S5f-j** and **Suppl. Table 6**). This is in line with many studies on astronauts and rodents flown at ISS^29,30^. In addition, our studies showed that spaceflight dramatically shifted mitochondrial metabolism toward more fatty acid and lipid oxidation, leading to high ROS production, mitochondrial deterioration, and global cellular damage^1^. IPA also revealed altered circadian rhythm in all five organs tested (**Suppl. Table 6**) and the dysregulation of circadian rhythm has a strong association with aging^31^. Given that evidence, our transcriptomic data argues for an altered aging process under spaceflight conditions.

Our integrative analysis provides novel insights into the tissue-specific and shared epigenomic and transcriptomic responses to spaceflight in mice. While global DNA methylation was relatively stable, spaceflight triggered widespread transcriptional reprogramming across organs, with consistent themes of mitochondrial dysfunction, immune suppression, and tissue-specific metabolic remodeling. These findings highlight key biological pathways that warrant further investigation and offer molecular targets for the development of countermeasures to safeguard astronaut health during long-duration missions. The dataset and analytical framework also offer a valuable resource for the broader space biology community and for comparative studies across future NASA and international spaceflight missions.

## Material and Methods

### Spaceflight and Mouse Tissue Collection

Mouse tissues used in this study were obtained from the NASA Biospecimen Sharing Program associated with the Rodent Research-18 (RR-18) mission, which was approved by the NASA Institutional Animal Care and Use Committee. Ten-week-old male C57BL/6 mice were sent to the International Space Station (ISS), where they were housed for 35 or 75 days. Corresponding ground control animals were maintained at the Roskamp Institute Animal Care Facility under identical housing conditions using NASA’s Animal Enclosure Modules (AEM). All mice were kept at an ambient temperature of 26–28 °C under a 12-hour light/dark cycle and were provided with the NASA Nutrient-upgraded Rodent Food Bar (NuRFB) and autoclaved deionized water ad libitum. Spaceflight animals were euthanized at three time points, along with their ground control counterparts, with a 3-day delay. Timepoint 1 (Frozen Carcass): One group of spaceflight mice was euthanized and immediately frozen on the ISS after 75 days in orbit. Ground control mice were euthanized and frozen simultaneously under matched conditions. Timepoint 2 (LAR_Acute): Another group of spaceflight mice was returned live to Earth aboard SpaceX’s Dragon capsule after 35 days in orbit and euthanized with 24 hours of splashdown. Timepoint 3 (LAR_3M): A final LAR group returned after 35 days on ISS was transported to Loma Linda University and was euthanized after a 3-month recovery period. All dissections were performed at the Loma Linda University Animal Care Facility by members of the RR-18 consortium. Organs and tissues were snap-frozen in liquid nitrogen and stored at –80 °C for long-term preservation.

### DNA/RNA Isolation and Library Construction

The genomic DNA (gDNA) and total RNA were isolated using the Qiagen AllPrep Universal Kit (Qiagen cat. 80224, Germantown, MD) according to the manufacturer’s protocol. gDNA and RNA were quantified using the Qubit dsDNA High Sensitivity Kit and RNA Broad Range Kit, respectively (Life Technologies, Carlsbad, CA). RNA quality was evaluated using the Agilent 2200 TapeStation and RNA ScreenTape (Santa Clara, CA). All RNA samples had an integrity number (RIN) above 8. The Purified RNA and DNA samples were kept at -80°C for long-term storage.

We processed 100 ng of isolated gDNA to generate RRBS DNA library using the EZ DNA Methylation Kit (Zymo D5001, Tustin, CA) according to the manufacturer’s protocol. Briefly, the methylation insensitive MspI enzyme, which cuts the DNA at CCGG sites, was used to digest gDNA into fragments. The fragments were directly subjected to end blunting and phosphorylation in preparation for ligation to a methylated adaptor. A unique index was used per sample for multiplexing. The ligation products were final repaired, then subjected to bisulfite conversion (98×C for 8 min, 54×C for 1h, hold at 4×C) using the lightning conversion reagent (Zymo D5030). Bisulfite-converted DNA was then amplified and purified using Zymo-Spin IC column.

Bulk RNA-seq libraries were constructed using Tecan Universal RNA-seq Library Preparation kit (Tecan, San Jose, CA) according to the manufacturer’s instructions. Briefly, 100 ng of total RNA was used to start cDNA synthesis. The products were then subject to end-repair, adaptor index ligation, and strand selection. A custom InDA-C primer mixture of SS5 Version 5 for mice was used to remove ribosomal RNA. Finally, libraries were amplified and purified with RNAClean XP Agencourt beads (Beckman Coulter, Indianapolis, IN); and quantified using Qubit dsDNA HS Kit on Qubit 4.0 Fluorometer (Life Technologies, Carlsbad, CA). The quality and peak size were determined using the D1000 ScreenTape on Agilent 2200 TapeStation (Agilent Technologies, Santa Clara, CA).

### RRBS and RNA-seq library sequencing

RRBS libraries were pooled and sequenced on Illumina NovaSeq X (UCLA Neuroscience Genomics Core) or DNBSEQ-G400GC sequencer (LLU Center for Genomics) at 2x150 paired-end reads. RNA-seq libraries were sequenced on Illumina NextSeq 550 (LLU Center for Genomics) at 75 bp single read.

### Mapping and processing of RRBS methylation data

The RRBS raw fastq files were first trimmed using Trim Galore (v0.4.5). Trimmed reads were aligned to the mouse reference genome NCBI GRCm38 with Bismark^32^ (v0.16.334) by default parameter settings. The methylation call files including the location of each CpG or CHH site and the methylation percentage were generated by the “bismark_methylation_extractor” function.

The output from bismark_methylation_extractor resulted in two files, one for CG methylation and another for CH (merged CHH and CHG) methylation. CG and CH methylation files were merged with coverage file to combine coverage and methylation percentage for each CG and CH. We obtained an average of 4.5 million mapped CpGs for each sample. We ensured that CpGs were detected in all samples in given tissues/organs. To generate consensus CpGs for each tissue, we ensured that CpGs were detected in all samples and required to have a minimum of 5 reads in at least 2 samples. Differentially methylated CpGs were identified using limma package^20^ with methylation change was >10% and p value of 0.0001.

### Mapping of RNA-seq data

For the RNA-seq data, we adopted the pipelines used in our recent publication^33^for RNA-seq data visualization, which integrated the QC (FastQC v0.11.4), trimming process (Trim_Galor v0.4.5), STAR alignment (v2.7.1a)^34^, reads quantification (HTseq v0.6.1)^35^ and differentially expressed gene (DEG) analysis (EdgeR v4.0.16)^36^. The genes with CPM ≥ 0.5 in all samples were used for DEG analysis. Gene count matrix was normalized using quasi-likehood (QL) dispersion. The DEGs were identified as |log2(Foldchange)| ≥ 0.5 and FDR :≤ 0.05.

### Enrichment analysis of Gene Ontology and KEGG pathways

For the enrichment of All DEGs identified in each organ/tissue were used for Gene Ontology (GO) functional analysis, using ShinyGo (v0.80, https://bioinformatics.sdstate.edu/go) and rat databases. FDR < 0.05 was used as cutoff to select enriched pathways. Enriched gene ontology were performed using clusterProfiler^21^.

### Enrichment of transcription factors motifs

Enrichment of motifs was performed using Homer^22^. For motif analysis in promoter regions, we extracted 1000 bases upstream and downstream of transcriptions start sites (TSSs) of differentially expressed genes. We performed de novo motif search in the promoter regions of differentially expressed genes in each tissue. Background sequences were generated by the Homer tool. After the first round of motif analysis, we selected significant (<1.0e-11) motifs from all tissues. We reran the motif analysis across significant motifs, which allowed to directly compare motifs across groups. Fold enrichment was calculated by measuring the motif signal in promoter versus background

## Supporting information

supplementary figures, and will be used for the link to the preprint site.

## Ethics Statement

All animal procedures were conducted in strict accordance with the National Institutes of Health (NIH) Guide for the Care and Use of Laboratory Animals and were approved by the NASA Institutional Animal Care and Use Committee (IACUC) as part of the Rodent Research-18 (RR-18) mission. The use of mice for spaceflight and ground control experiments, including housing, handling, euthanasia, and tissue collection, was reviewed and approved under NASA’s Biospecimen Sharing Program. Ground control animals were maintained under conditions matched to spaceflight animals using NASA Animal Enclosure Modules (AEM). All post-flight animal handling, dissections, and tissue processing were performed at the Loma Linda University Animal Care Facility by trained personnel from the RR-18 consortium, in compliance with institutional and federal animal welfare regulations. All efforts were made to minimize animal suffering and to reduce the number of animals used. The ethics approval number is LLU IACUC# 8170051.

## Authors’ contributions

CW conceived, designed the study, and provided funding. VM, MP, and MB provided some spaceflight animal tissues for some organs. ZC and FZ performed the experiments. ZC and CN performed bioinformatics data analysis and drafted the manuscript. ZC, CN, and CW revised the manuscript. All authors reviewed and agreed with the final version of the manuscript.

## Acknowledgements

The authors would like to thank the NASA RR-18 Biospecimen Sharing Program for providing the biological material, with special thanks to America Reyes Wang from NASA’s Ames Research Center for leading the program. We would like to thank Ms. Adriana Lopez of the LLU Center for Genomics for her administrative support and Dr. Jacob Holley for his technical assistance with the RRBS library preparation. This study was supported by NASA Space Biology grant # NNX15AB41G. The genomic work carried out at the LLU Center for Genomics was funded in part by the National Institutes of Health (NIH) grants S10OD019960 (CW), U01DA058278 (CW), the Ardmore Institute of Health grant 2150141 (CW) and Dr. Charles A. Sims’ gift to LLU Center for Genomics.

