## supplementary figures, and will be used for the link to the preprint site. for "A Comprehensive DNA Methylome BodyMap across 12 Organs/Tissues from Spaceflight Mice"

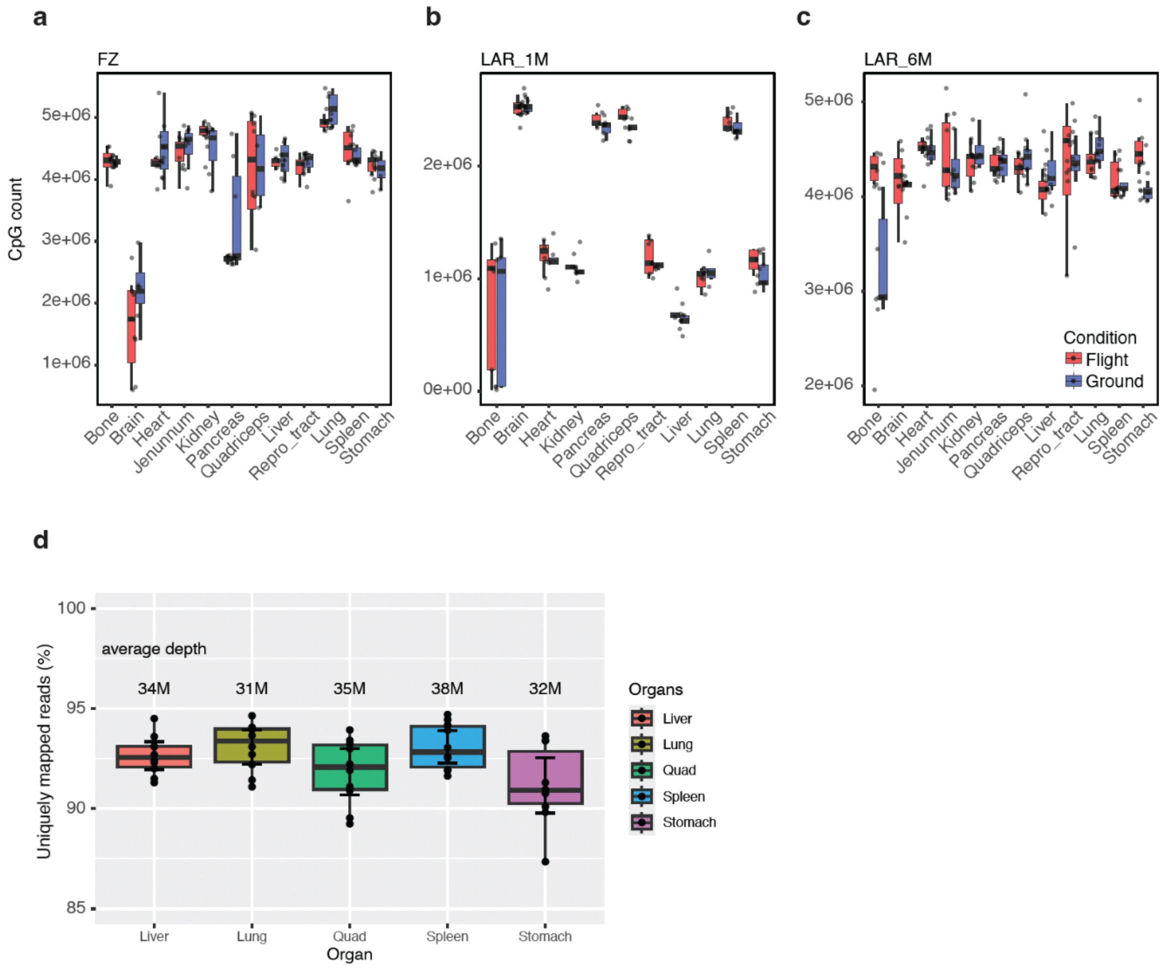


**Supplemental Figure 1. Mapped CpGs and RNA-seq reads across tissues/organs and experimental conditions.** (a-c) Boxplots showing the number of CpGs mapped per sample in ground and spaceflight conditions across three experimental setups (a) spaceflight frozen (FZ) samples, (b) live animal return (LAR) with 30 days recovery (LAR_1M) and (c) live animal return (LAR) with 6 months recovery (LAR_6M). (d) Boxplots showing RNA-seq read depth and mapping efficiency across different organs.

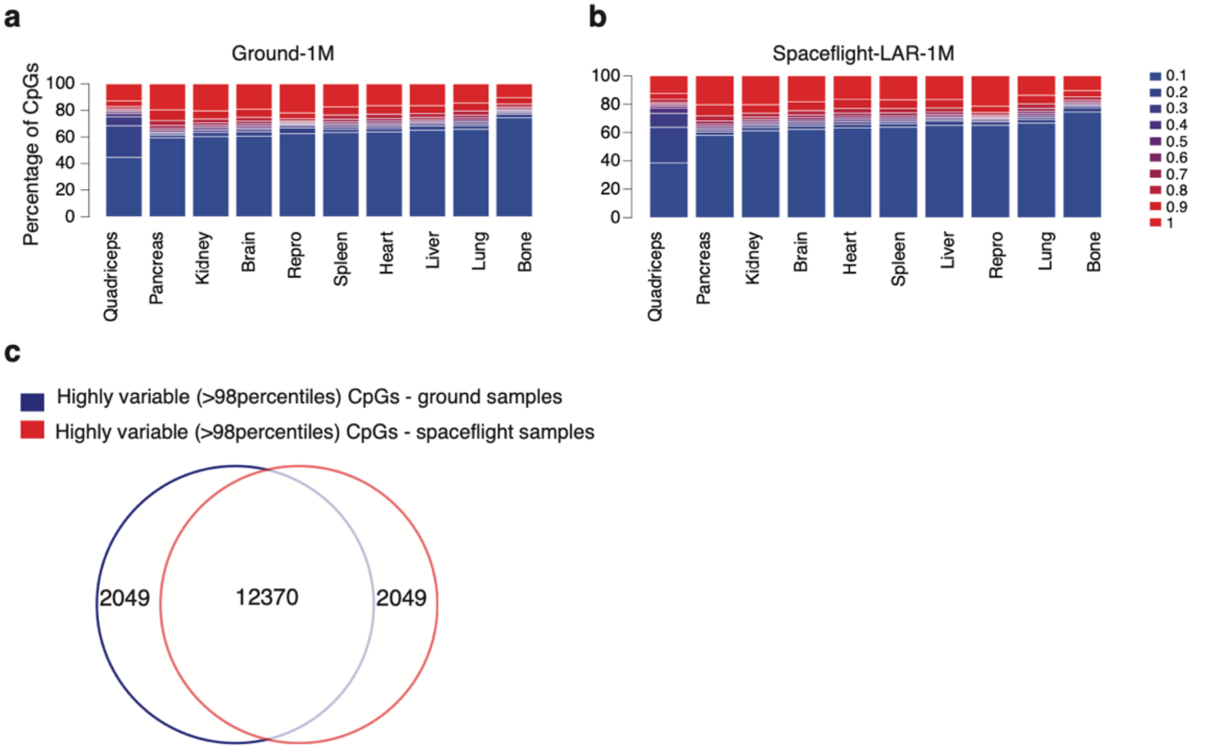


**Supplemental Figure 2. DNA methylation landscape across tissues/organs and experimental conditions. (a-b)** Distribution of mean beta values of all CpGs across all tissues in (a) ground control and (b) spaceflight conditions. Beta values are divided into ten bins. Y-axis represents the percentage of CpGs per bin. (**c**) A Venn diagram showing the overlap between highly variable (>98th percentiles) CpGs across all tissues from ground and spaceflight conditions.

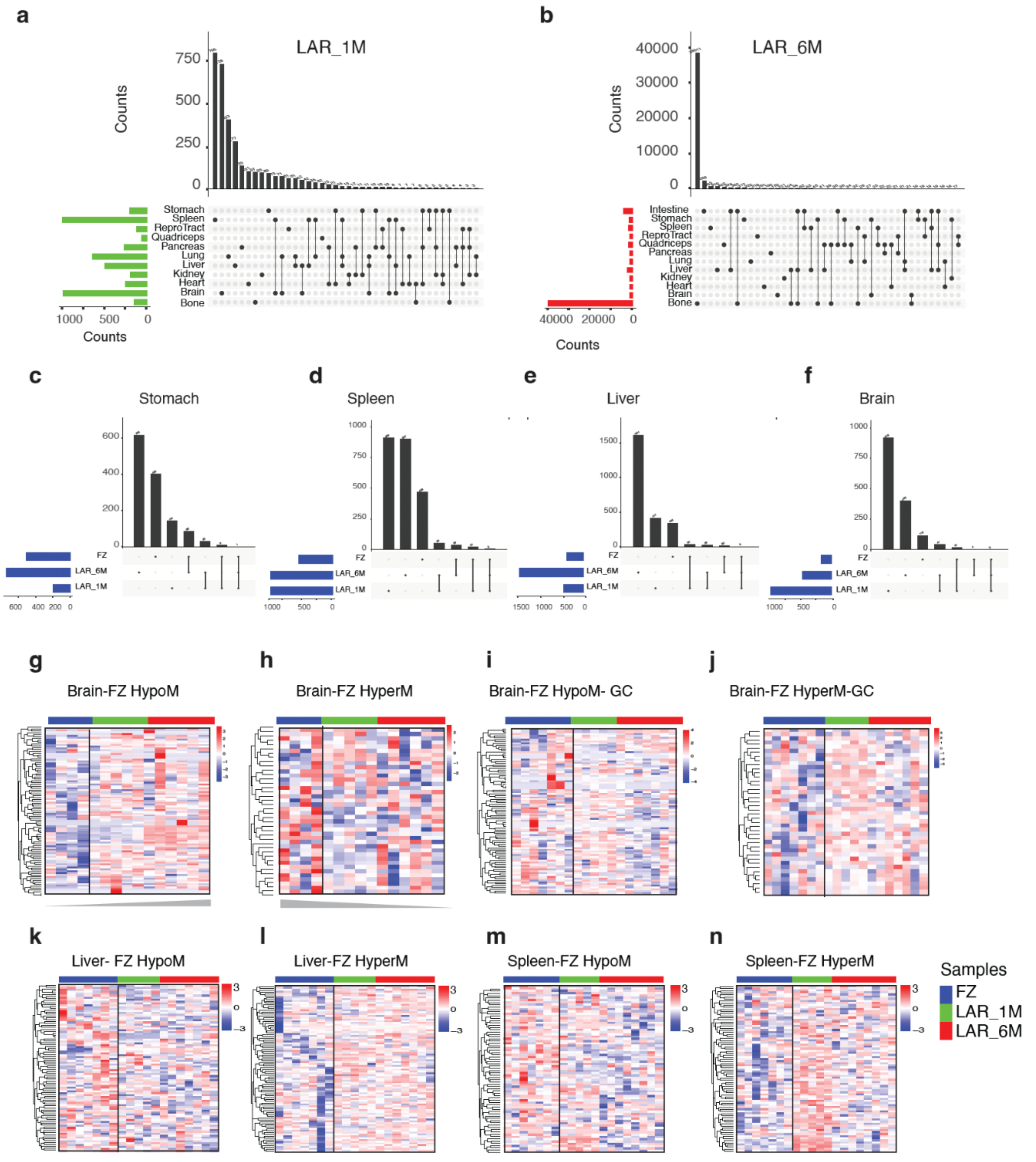


**Supplementary Figure 3. Methylation landscape of differentially methylated CpGs (DMCs) in liver animal return (LAR) recovered for 1 month (LAR_1M) and 6 months (LAR_6M) groups.** (**a-b**) Overlap of DMCs between different organs/tissues in (a) LAR_1M and (b) LAR_6M groups. (c-f) Overlap of differentially methylated CpGs from (c) stomach, (d) spleen, (e) liver, (f) brain tissues during flight (FZ) and two recovery time points (LAR_1M and LAR_6_M). **(g-h**) Heatmaps of brain hypomethylated CpGs (g) and hypermethylated CpGs (h) identified in flight (FZ) samples and their methylation levels in corresponding recovery samples at LAR_1M and LAR_6M. Methylation values are row-scaled. (**i-n**) Heat maps show the methylation levels of differential CpGs across in spaceflight samples of brain, liver and spleen in their corresponding ground control across three time points (FZ, LAR_1M and LAR_6M).

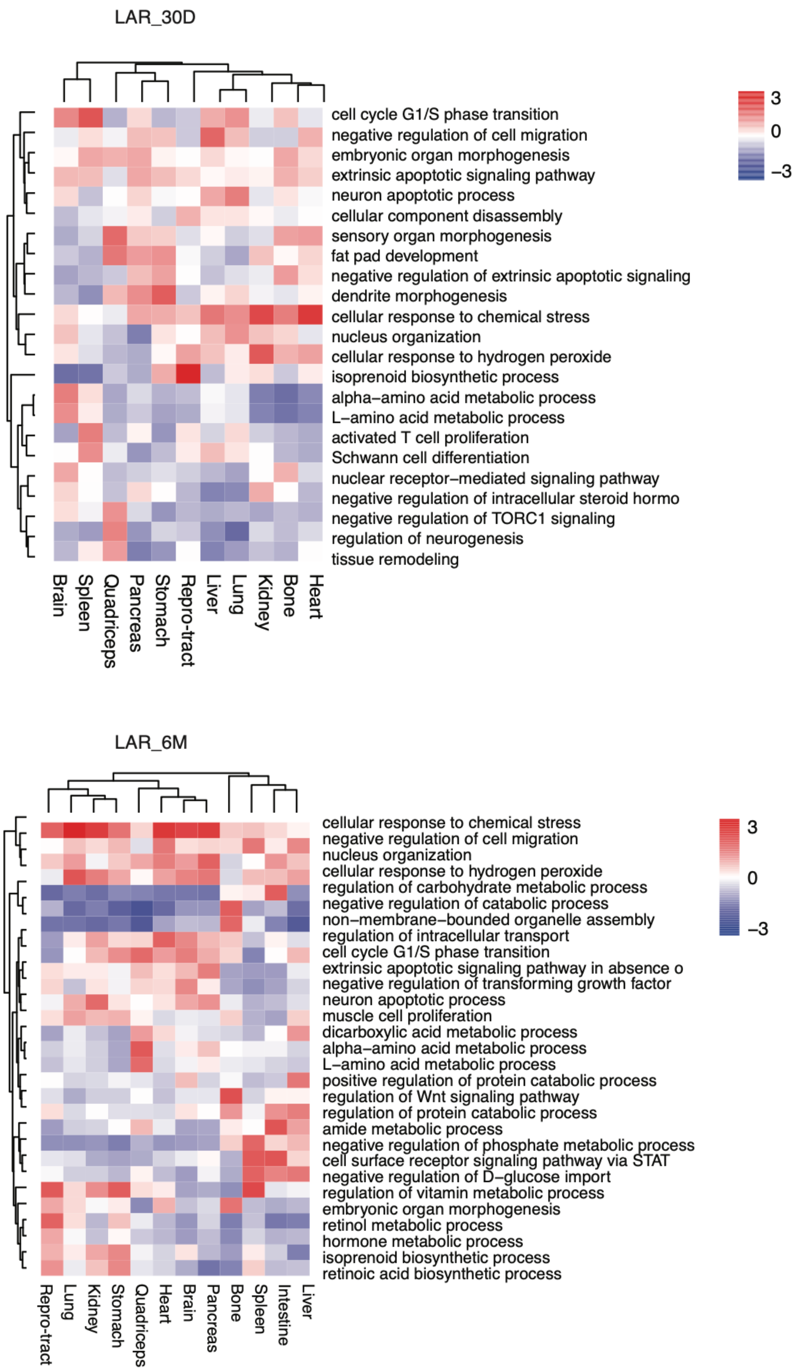


**Supplementary Figure 4. Enrichment of gene ontology associated with differentially methylated CpGs (DMCs) from two recovery groups, live animal return (LAR) recovered for one month (LAR_1M) and six months (LAR_6M).** (a) Heatmap shows enriched gene ontology terms on genes proximal to DMCs from LAR_1M group. P-values are scaled across rows. (b) Heatmap shows enriched gene ontology terms on genes proximal to DMCs from LAR_6M group. P-values are scaled across rows.

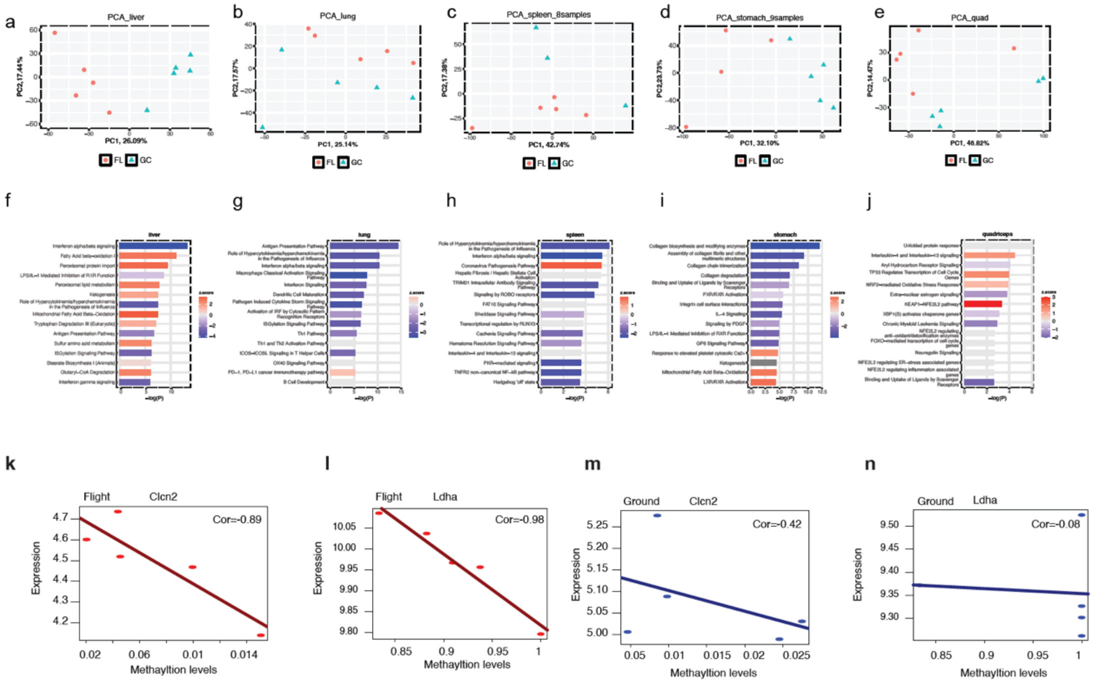


**Supplementary Figure 5. Spaceflight-induced differential transcriptomes across multiple mouse organs. (a-e)** Principal component analysis (PCA) plots showing spaceflight-induced transcriptomic changes in liver (a), lung (b), spleen (c), stomach (d), and quadriceps (e). **(f-j)** Bar-plots showing the IPA canonical pathways enriched by the spaceflight induced DEGs in liver (f), lung (g), spleen (h), stomach (i), and quadriceps (j). **(k-n)** Correlation of gene expression and DNA methylation levels of genes Clnc2 (k and m) and Ldha (l and n) in flight and ground conditions.

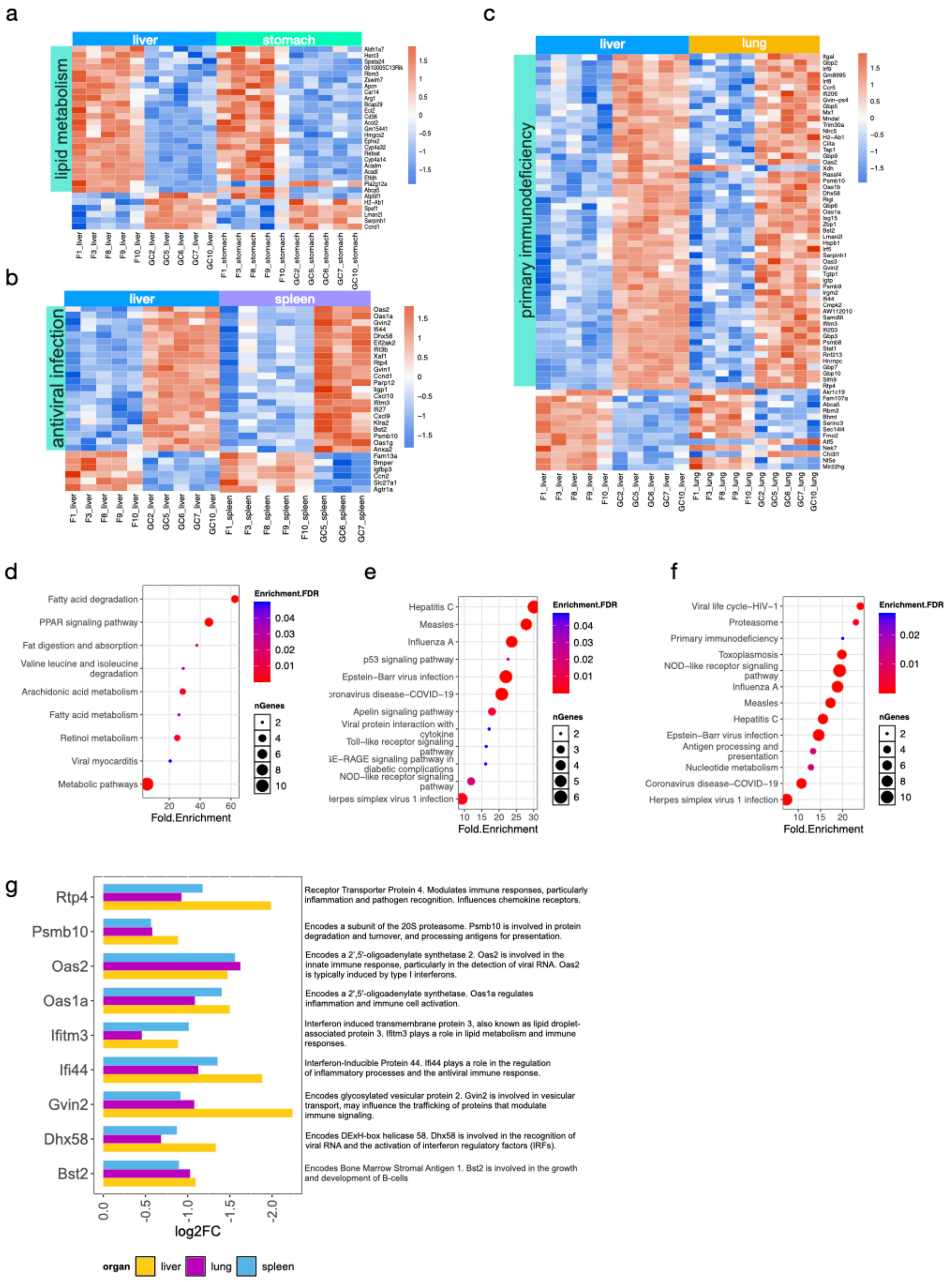


**Supplementary Figure 6. Synergistic transcriptomic responses of rat organs to spaceflight.** Heatmaps showing the expressions of the common DEGs between liver and stomach (a), liver and spleen (b), and liver and lung (c). The expression was scaled in row level. The biological functions of the DEGs were labelled on the left of the heatmaps. Gene ontology biological processes enriched from the common DEGs between liver and stomach (d), liver and spleen (e), and liver and lung (f). (g) The relative expression and functions of the nine common DEGs identified in liver, lung, and spleen.
